# An Optically Decodable Bead Array for Linking Imaging and Sequencing with Single-Cell Resolution

**DOI:** 10.1101/355677

**Authors:** Jinzhou Yuan, Jenny Sheng, Peter A. Sims

**Affiliations:** Department of Systems Biology, Columbia University Medical Center, New York, NY 10032 USA; Integrated Program in Cellular, Molecular, and Biomedical Studies, Columbia University Medical Center, New York, NY 10032 USA; Department of Biochemistry & Molecular Biophysics, Columbia University Medical Center, New York, NY 10032 USA; Sulzberger Columbia Genome Center, Columbia University Medical Center, New York, NY 10032 USA

**Keywords:** single-cell RNA-Seq, live cell imaging, microfluidics

## Abstract

Optically decodable beads link the identity of an analyte or sample to a measurement through an optical barcode, enabling libraries of biomolecules to be captured on beads in solution and decoded by fluorescence. This approach has been foundational to microarray, sequencing, and flow-based expression profiling technologies. We have combined microfluidics with optically decodable beads to link phenotypic analysis of living cells to sequencing. As a proof-of-concept, we applied this to demonstrate an accurate and scalable tool for connecting live cell imaging to single-cell RNA-Seq called Single Cell Optical Phenotyping and Expression (SCOPE-Seq).

## Main Text

Recent advances in microfluidics and cDNA barcoding have led to a dramatic increase in the throughput of single-cell RNA-Seq (scRNA-Seq)[1-5]. However, unlike earlier or less scalable techniques[6-8], these new tools do not offer a straightforward way to directly link phenotypic information obtained from individual, live cells to their expression profiles. Nonetheless, microwell-based implementations of scRNA-Seq are compatible with a wide variety of phenotypic measurements including live cell imaging, immunofluorescence, and protein secretion assays[3, 9-12]. These methods involve co-encapsulation of individual cells and barcoded mRNA capture beads in arrays of microfabricated chambers. Because the barcoded beads are randomly distributed into microwells, one cannot directly link phenotypes measured in the microwells to their corresponding expression profiles. In Single Cell Optical Phenotyping and Expression Sequencing or SCOPE-Seq, we use optically barcoded beads[13] and identify the sequencing barcode associated with each single-cell cDNA library on the sequencer by fluorescence microscopy. Thus, we can obtain images, movies, or other phenotypic data from individual cells by microscopy and directly link this information to genome-wide expression profiles.

We previously demonstrated the compatibility of the commercially available “Drop-seq” beads[1] with our microwell array system for scRNA-Seq[9, 14]. Here we generate optically barcoded mRNA capture beads for our microwell array system from “Drop-seq” beads. These beads are conjugated to oligonucleotides with a cell-identifying sequencing barcode and a 3’-poly(dT) terminus. To enable optical identification of the sequencing barcode on each bead, we attach a unique combination of oligonucleotides selected from a set of 12 in two cycles of split-pool ligation (**Fig. 1A**, left). We refer to this combination as an optical barcode, because it can be decoded by sequential fluorescence hybridization, to distinguish it from the sequencing barcode. Immediately prior to ligation, we sonicate the beads to detach a small fraction of oligonucleotides, from which we generate a “bead-free” sequencing library (Supplementary Methods, **Fig. 1A**, right). Split-pool, single-stranded ligation produces a final pool of beads with both optical and sequencing barcodes (dual-barcoded beads, **Fig. 1B**). Sequencing the bead-free library associated with each ligation reaction produces a look-up table linking the optical and sequencing barcodes for each bead in the pool (**Fig. 1C**). Importantly, bead-free library construction and sequencing is done once for each batch of optically barcoded beads, which can be used for many SCOPE-Seq experiments.

**Figure 1.**
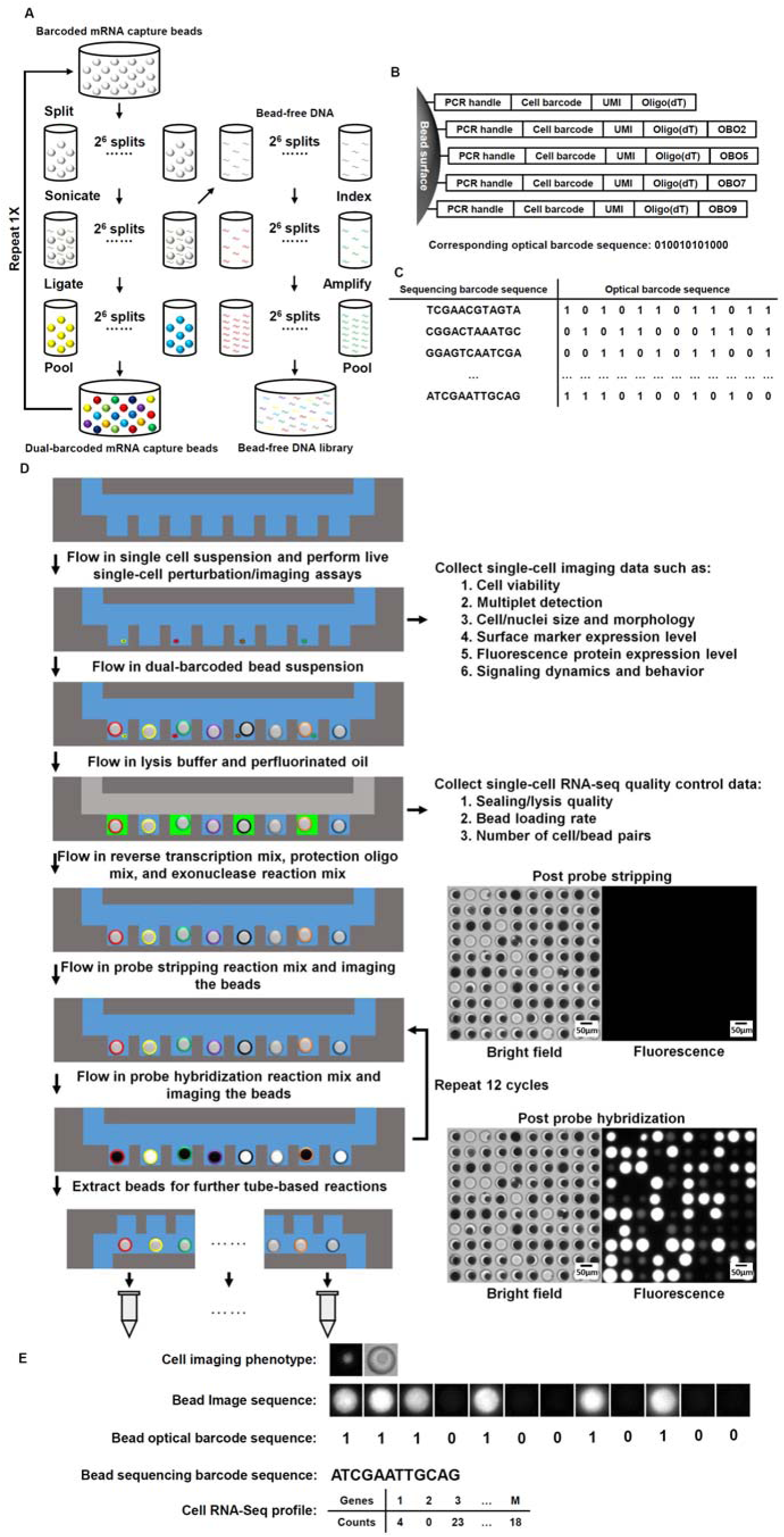
A) Workflow for generating the dual-barcoded beads and the associated bead-free DNA sequencing libraries. B) Sequence composition of the bead-bound mRNA capture oligonucleotides and optical barcode oligonucleotides (OBOs). C) A look-up table linking sequence barcodes and optical barcodes. D) Workflow for SCOPE-Seq. E) Workflow for linking live, single-cell imaging data with single-cell RNA-Seq profile.

In SCOPE-Seq, we load cells into a microwell array as described previously[3, 9], collect live cell imaging data, and co-encapsulate the cells with dual-barcoded beads (**Fig. 1D**). The microwell array device used here has a total of 30,500 microwells. We then use a computer-controlled system to perform on-chip cell lysis, mRNA capture, reverse transcription, and exonuclease digestion. This process generates PCR-amplifiable, sequence barcoded cDNA that is covalently attached to each bead. Next, we perform “optical demultiplexing” - 12 cycles of reversible fluorescence hybridization that determine the combination of 12 optical barcode oligonucleotides (OBOs) on each bead (Supplementary Methods, **Fig. 1D-E**). We then cut the microwell array into multiple pieces and extract the beads from each piece for scRNA-Seq library construction. Beads from each piece are processed and indexed separately, thereby increasing our multiplexing capacity to 2^12^\**N* where N = 10 is the number of pieces, giving us an effective barcode library size of 40,960. The resulting scRNA-Seq libraries from different pieces are then pooled and sequenced to obtain the RNA-Seq profile of the individual cells and their sequencing barcodes. Finally, we process the images to identify the optical barcode on each bead and identify the corresponding sequencing barcode using the look-up table described above to link microscopy data from a cell in a particular microwell to its RNA-Seq profile (**Fig. 1E**).

To characterize the performance and demonstrate the utility of SCOPE-Seq, we performed a mixed-species experiment where human (U87) and mouse (3T3) cells are labeled with differently colored live stain dyes, mixed together, and analyzed with SCOPE-Seq. Differential labeling allows us to determine the species of each captured cell by live cell imaging in the microwells. We can independently determine the species associated with each sequencing barcode using the corresponding scRNA-Seq data, from which we detected ~3,600 unique transcripts per cell on average. We considered any inconsistencies between these two independent species identifications to result from optical barcode linkage error. We define the linking accuracy as the fraction of single-cell imaging and RNA-Seq data sets that are correctly linked among all linked data sets. **Fig. 2A** shows a scatter plot of the number of reads aligning uniquely to the human and murine transcriptomes for each cell that was linked to imaging data. Importantly, the data points derived from scRNA-Seq are colored using the actual two-channel fluorescence data from live cell microscopy. Species calling was 98.6% concordant between the imaging and scRNA-Seq data, and after correcting for the difference in abundance of human and murine cells, we obtained an imaging-to-sequencing linking accuracy of 96.1% for singlets from which we can unambiguously obtain a species call. From a total of 2,352 RNA-Seq expression profiles, 1,133 of them were linked to imaging data (48.2%). The majority of unlinked cells are due to optical demultiplexing errors which usually lead to a failure to link rather than an error. Because the yield of linked optical and sequencing barcodes is high (89.6%), we know that these errors arise from the optical demultiplexing process itself.

**Figure 2.**
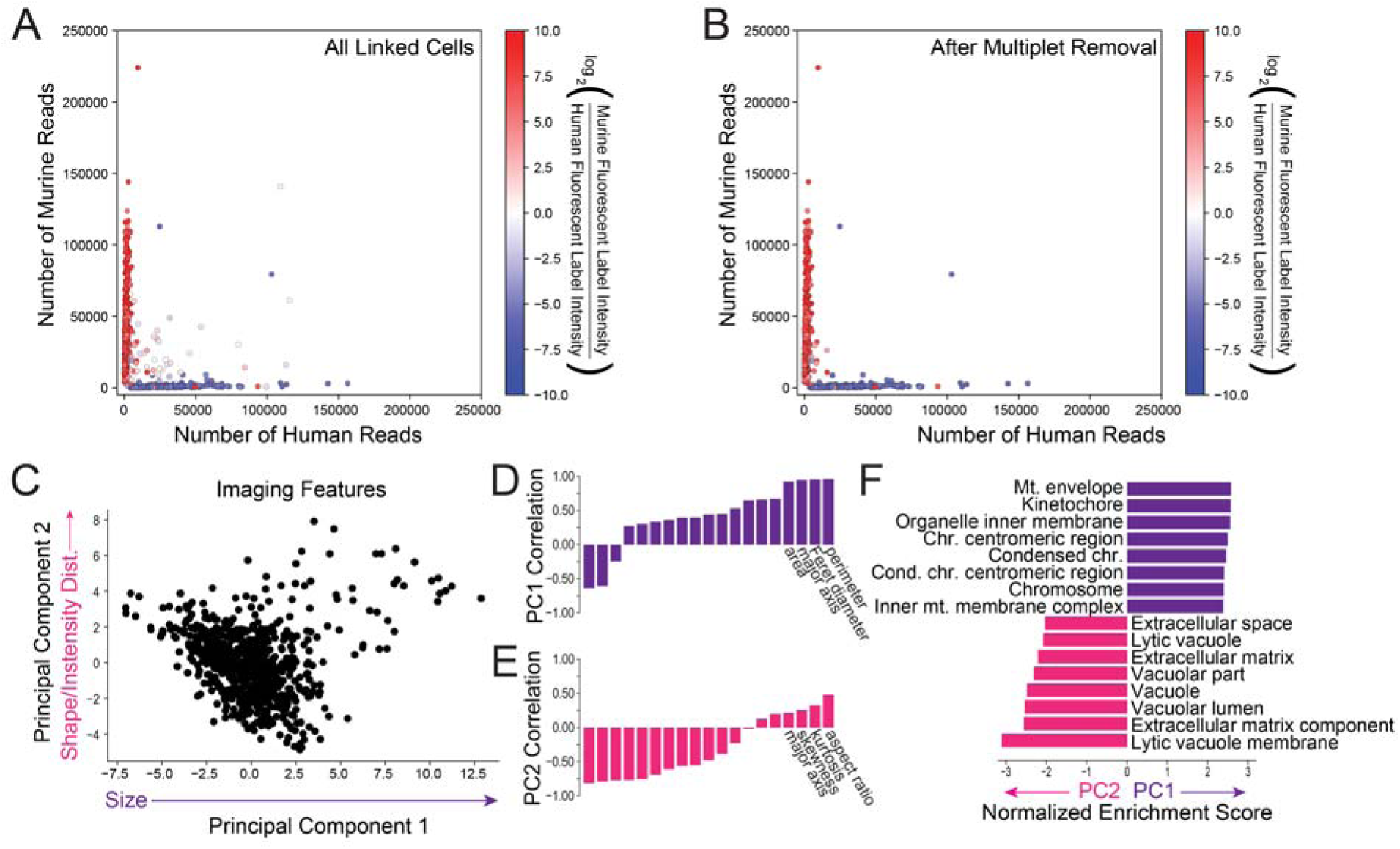
A) Scatter plot of the number of uniquely aligned human and murine reads for each linked sequencing barcode colored by the relative fluorescence intensity of the human and murine labels before and B) after imaging-based multiplet removal. C) Principal component analysis of the 19 fluorescence imaging features for the murine cells. D) Correlation between the first PC and each imaging feature. E) Correlation between the second PC and each imaging feature. F) Normalized enrichment scores for MSigDB gene sets enriched in genes that are correlated with the first (purple) and second (magenta) PCs.

The microwell array contains a small number of multi-species multiplets (**Fig. 2A**). As expected from the high linking accuracy of SCOPE-Seq, the species purity of RNA-Seq expression profiles are consistent with their fluorescent labels from live-cell imaging data (**Fig. 2A**). Interestingly, we were able to remove most of the multi-species multiplets using monochrome live-cell imaging data (Supplementary Methods, **Fig. 2B**). We used scRNA-Seq profiles with purity below 70% as a threshold for calling human-mouse multiplets. We then blinded ourselves to the two-color fluorescence information that distinguishes human-mouse multiplets in our imaging data and attempted to identify multiplets by manually examining the monochrome images. Imaging- and sequencing-based doublet identifications were conducted by different researchers in a blinded fashion. This resulted in a sensitivity of 66% and a specificity of 99.1% for multiplet detection, and a concordance of 87.5% between one- and two-color imaging. Some false negatives likely arise from imperfections in our scRNA-Seq “ground truth” and relatively low-resolution imaging. We anticipate that more sophisticated image processing and better microscopy and cell/nucleus stains will lead to further improvements in sensitivity to make SCOPE-Seq highly effective for detecting multiplets.

Additional imaging features obtained from cells may carry information related to gene expression. We measured 19 imaging features from the fluorescence images of cells reflecting various aspects of cell size, shape, and intensity distribution (Supplementary Methods). Principal component (PC) analysis on these features suggests significant heterogeneity among the cells (**Fig. 2C**). The first PC is primarily correlated with cell size-related features (**Fig. 2D**) and the second with shape-related features (**Fig. 2E**). We then ranked protein coding genes by their correlation with these PCs and performed gene set enrichment analysis for each PC (**Fig. 2F**). Perhaps not surprisingly, we found that cell division-related genes are most correlated with the first PC, which is related to cell size, likely because actively dividing cells and mitotic figures tend to be larger. We found enrichment of extracellular matrix vacuolar genes for the second PC, which represents a measure of cell shape and may result from cells in the process of adhering to the microwells and the impact of vacuoles on overall cell morphology. We made very similar observations of associations between gene ontology and imaging features using partial least squares regression analysis on the same data (Supplementary Methods, **Supplementary Table 2**).

Many cellular phenotypes are difficult to infer directly from static measurements of the transcriptome, such as protein expression and localization dynamics, organelle dynamics and distribution, morphological features, uptake of foreign objects, and biomolecular secretion. However, there are myriad live cell imaging and microwell-based assays for characterizing these phenotypes in individual cells. We expect that SCOPE-Seq will serve as a highly scalable, accurate, and economical approach to linking live cell microscopy assays to scRNA-Seq, enabling investigation of the transcriptional underpinnings of the resulting phenotypes. In addition, with the multiplet detection capability, one can potentially achieve ~5-fold higher throughput by more aggressive cell loading, although this is subject to the caveat that background from molecular cross-talk could increase.

The current implementation of SCOPE-Seq has important limitations, motivating future improvements. The use of separate optical and sequencing barcodes requires an extra step-sequencing of bead-free libraries. Future implementations will use oligonucleotides in which the optical and sequencing barcodes are the same DNA sequence. OBO ligation to a subset of mRNA capture sites on the beads also leads to reduced molecular capture efficiency for the corresponding scRNA-Seq libraries relative to beads without optical barcodes (**Supplementary Fig. 11**). Finally, the multiplexing capacity of the beads in our proof-of-concept experiment is relatively modest, which precludes an error-correcting code. While our current linking accuracy is high, both the yield of optically demultiplexed cells and accuracy could be improved with such a code.

The scheme presented here for linking cellular phenotypes to sequencing could also be generalized beyond single-cell analysis. Microscopy experiments on small multi-cellular organisms, organoids, or colonies could be linked with sequencing. Additionally, the optically decodable bead array could be used for spatial transcriptomic analysis as demonstrated previously with printed microarrays[15]. One advantage of optically decodable beads over printed microarrays[15, 16] is that beads can be prepared in a large batch for use in many experiments starting from commercially available “Drop-seq” beads with relatively simple tube-based reactions. We hope that this economical approach will serve as a powerful tool for connecting high-throughput microscopy and sequencing on multiple scales.

## Availability of Data and Materials

The single-cell RNA-Seq data and bead-free optical barcode/sequencing barcode linkage data have been deposited in the Gene Expression Omnibus under accession GSE116011.

## Author Contributions

J.Y. and P.A.S. conceived the study and designed experiments. J.Y. constructed the automated instrumentation for SCOPE-seq and conducted all of the experiments. J.Y., J.S., and P.A.S. analyzed the data. J.Y. and P.A.S. wrote the paper.

## Acknowledgements

We thank the Sulzberger Columbia Genome Center for assistance and resources for high-throughput sequencing.

